# CAPG is required for Ebola virus infection by controlling virus egress from infected cells

**DOI:** 10.1101/2022.04.10.487814

**Authors:** Hiroyuki Mori, James P Connell, Callie J Donahue, RuthMabel Boytz, Justin J Patten, Daisy Leung, Douglas J LaCount, Robert A Davey

**Affiliations:** Department of Microbiology, NEIDL, Boston University School of Medicine, Boston, MA, USA; Department of Medicinal Chemistry and Molecular Pharmacology, Purdue University, West Lafayette, IN, USA; Division of infectious diseases, Washington University School of Medicine, St. Louis, MO, USA

**Keywords:** Ebola virus, filovirus, CAPG, actin, virus assembly, host factors

## Abstract

Replication of Ebola virus (EBOV) is dependent upon actin functionality, especially at cell entry through macropinocytosis and at release of virus from cells. Previously, major actin-regulatory factors such as Rac1 and ARP2/3, involved in actin nucleation were shown important in both steps. However, downstream of nucleation, many other cell factors, are needed to control actin dynamics. How these regulate EBOV infection remains largely unknown. Here, we identified the actin-regulating protein, CAPG, as important for EBOV replication. Notably, knockdown (KD) of CAPG specifically inhibited viral infectivity and yield of infectious particles. Mechanistic analysis revealed a requirement of CAPG for virus production from infected cells. Proximity ligation and split GFP reconstitution assays revealed strong association of CAPG with VP40 that was mediated through the S1 domain of CAPG. Overall, CAPG is a novel host factor regulating EBOV infection through connecting actin filament stabilization to viral egress from cells.

## Introduction

EBOV is a cause of severe hemorrhagic disease with a high mortality rate in humans and nonhuman primates [1,2]. Sporadic outbreaks, including the 2013-2016 outbreak in Western Africa, have threatened human health and society significantly, and EBOV continues to be a global health burden [3,4]. Even though much molecular and clinical research has been conducted to develop promising therapeutics, only a limited number of treatments and vaccines have been approved by the FDA [4–6]. These therapeutics must be administered early in disease onset to be effective [7] and are difficult to deploy in outbreak conditions [8]. The limited availability of these countermeasures as well as the risk of viral resistance accentuates the need for further therapeutic development[6]. Since the virus life cycle depends on the host cell and cellular machinery, host factors provide attractive targets for alternative antiviral drug development[9]. Thus, understanding the molecular mechanisms of EBOV infection in host cells and the subsequent interactions of host and virus factors is necessary for combating future outbreaks and disease.

EBOV is a non-segmented, negative-sense RNA virus, belonging to the family Filoviridae [10]. Virions are morphologically characterized by diameters of ~80 nm and lengths ranging from hundreds of nanometers to micrometers [11,12]. The filovirus genome encodes seven viral proteins, all of which are present in intact virions [13]. The viral nucleocapsid is composed of the RNA genome in association with the nucleoprotein (NP), the viral polymerase L, and replication and transcription factors VP35 and VP30 [14,15]. VP40, the viral matrix protein, contains a PPXY motif, capable of interacting with a variety of host cell proteins [16] and is the most abundant protein in the virion [13]. During late stages of virus assembly, VP40 connects the viral nucleocapsid to the host-membrane derived viral envelope [16–18], which is studded with the viral glycoprotein GP [12], a type 1 transmembrane protein composed of trimeric post-translationally cleaved GP subunits GP1 and GP2 [19–21]. GP serves as the major mediator of viral entry into cells [22,23].

EBOV enters cells via macropinocytosis [24,25], a cellular process heavily dependent on actin function [26,27]. Macropinocytosis is driven by membrane ruffling which is mediated by cortical actin branching at the cytosolic surface of the plasma membrane [26,27]. This actin remodeling is dependent on the actions of Rho-GTPases such as Rac1 and Cdc42, in addition to the multimeric protein complex Arp2/3 [26,27], all of which are regulated upstream by PI3 kinase (PI3K) signaling pathway[26]. We and others previously showed that inhibition of PI3K and Rac1 interferes with cellular uptake and entry of EBOV [24,28] and additional studies implicated the involvement of additional Rho – GTPases in EBOV entry [29], ultimately demonstrating a heavy involvement of actin dynamics in successful viral entry and overall infection.

Actin polymerization has been additionally shown to participate in late stages of EBOV infection, notably in assembly and egress. [11,30–33]. Assembly initiates with the formation of the nucleocapsid, a process driven by NP, VP35, and VP24 [33–37]. VP24 facilitates viral transcription and replication and ensures packaging of the viral genome into the nucleocapsid [11,35–39]. The nucleocapsid is then transited to sites of VP40-driven viral budding at the plasma membrane via actin polymerization [32,40], a process that is also dependent on actin nucleation by ARP2/3 [32,41]. VP40 is capable of inducing membrane curvature and can independently induce viral budding [17,31], though budding is enhanced with expression of GP [18,42]. Through interaction with host factors TSG101 and VPS4, VP40 coordinates release of viral particles [43,44]. Like nucleocapsid transit, VP40-driven budding is also dependent on actin-functionality [30,45–47] with actin has been detected within VP40 budding protrusions and virus-like particles (VLPs) [30]. Inhibitors of calmodulin, a calcium sensing protein capable of regulating filamentous actin (F-actin) formation in cells [48] similarly inhibits VP40 driven budding and egress [49]. Successful production of infectious virions therefore is intimately connected with actin processing and production.

Actin remodeling is controlled by interaction with actin-interacting proteins such as cofilin, gelsolin and fascins, through actin monomer polymerization to F-actin, depolymerization and branching [50–54]. Macrophage-capping protein (CAPG) is one such protein that is a member of the Gelsolin/Villin family which binds to the barbed-end of F-actin and responds to Ca^2+^ and phosphoinositide signals (**Fig. S1A**). The main function of the family is to promote stability of the actin filament by capping the growth end of the filament to prevent further elongation while preventing loss of actin subunits [55]. CAPG is unique among its family members, consisting of only three gelsolin homology (GH) domains while others have six and lacks actin severing activity [56]. In addition, CAPG is observed both in the nucleus and in the cytoplasm, while gelsolin-like proteins localize predominantly in the cytoplasm [57,58]. The role of CAPG and the Gelsolin/Villin family in macropinocytosis, as used by EBOV to enter cells, is not fully understood. However, it has been shown that CAPG and gelsolin double-null mice exhibit diminished membrane ruffling and protrusion [59], which are characteristics of macropinocytosis [27].

In this study, we studied the role of CAPG on EBOV infection for cell entry, replication, and budding from infected cells. We show that CAPG plays only a minor role in virus uptake into cells. Instead, it plays a predominant role in virus egress from cells. This appears to be mediated by close association with the major nucleocapsid protein, VP40.

## Materials and Methods

### Cells and culture

Human cervical carcinoma (HeLa) cells and Vero E6 cells (ATCC, Manassas, VA) were cultivated in Dulbecco’s modified Eagle’s medium (DMEM) (11995073; Gibco, Gaithersburg, MD) supplemented with 10% fetal bovine serum (FBS) (S10350; R&D systems, Minneapolis, MN) and 1% penicillin-streptomycin solution (15140-122; Gibco). All cells were maintained at 37°C in a humidified incubator with 5% CO_2_.

### Reagents and antibodies

Hoechst 33342 dye (62249) and HCS CellMask blue (H32720) for immunostaining were purchased from ThermoFisher Scientific (Waltham, MA). Mouse monoclonal anti-ZEBOV GP (4F3) and anti-ZEBOV VP40 (3G5) antibodies were obtained from IBT Bioservices (Gaithersburg, MD). Mouse monoclonal anti-ZEBOV NP (10B5 C11 C9 C3), and VP35 (2A11 F12 D7) were obtained by Dr. D. Leung (Wash. U. St. Louis, MI). Rabbit polyclonal anti-Ebola VP30 (GTX134035) was purchased from GeneTex (Irvine, CA). Rabbit polyclonal anti-CAPG antibody (10194-1-AP) was from Proteintech (Rosemont, IL). Mouse monoclonal anti-*β*-actin (MAB8929) and rabbit polyclonal anti -HSP60 (AF1800) antibodies were purchased from R&D systems. The following secondary antibodies were purchased from Thermo Scientific: goat anti-mouse Alexa Flour 488, anti-mouse Alexa Flour 546, anti-rabbit Alexa Flour 488 and anti-rabbit Alexa Flour 546 5-(N-ethyl-N-isopropyl) amiloride (EIPA) and chlorpromazine hydrochloride (CPZ) were from Sigma-Aldrich (St. Louis, MO). EIPA was diluted to 10 mM with dimethylsulfoxide. CPZ was diluted to 50 mg/ml with water. All the reagents described here were kept in −20°C until use.

### Plasmids

All plasmids indicated were purified by either PureLinkTM HiPure Plamid Midiprep kit (K210004; ThermoFisher Scientific) or Plasmid Maxi kit (12162; Qiagen). pCAGGS-VP40, pCAGGS-eGFP-VP40, pcDNA3-GP, pC-NP, pC-VP30, pC-VP35, and pC-L constructs are described elsewhere [24,60–62]. Each plasmid expresses the Mayinga variant of the indicated EBOV protein. For the tripartite split-GFP system, GFP B10 and B11 fusion constructs were created in pCDNA3- and pCDNA5-based plasmids (PMID: 24092409). GFP B10 was fused to EBOV VP40 and VP30 at the 5’ and 3’ ends, respectively, whereas GFP B11 was fused to human genes or gene fragments (5’ fusions for CAPG, TSG101, and utrophin CH domain; 3’ fusions for CAPG and PABPC1). Genes were PCR-amplified and inserted by ligation or NEBuilder HiFi DNA Assembly (New England Biolabs, E2621) or transferred from another plasmid by restriction digest and ligation. All plasmids were confirmed by DNA sequencing. Plasmid maps and sequences can be found in supplemental material. Primer sequences and cloning details are available upon request. PCDNA3-GFP1-9 T2A mCherry was a gift from Xiaokun Shu (Addgene plasmid # 124430;http://n2t.net/addgene:124430; RRID:Addgene_124430) (PMID: 30821975). Additional details can be found in the **Supplementary Materials and Methods**.

All infection assays were performed in triplicate. HeLa cells were seeded in 96, 24, 12 or 6 well plates (3596, 3524, 3512, 3506, respectively; Corning, Glendale, AZ) and incubated overnight at 37°C. Only wells showing 50-70% confluency of cells were used for subsequent infection assays. The cells were then challenged with WT-EBOV or EBOV-GFP and incubated for 1 h at 37°C. The supernatant was removed, and cells were washed twice with DMEM without serum. Fresh DMEM supplemented with 10% FBS was added, and cells were incubated for the appropriate time depending on the assay to be performed. The MOI for all infections was 0.2 unless otherwise indicated. All experiments with replication-competent EBOV were performed in a biosafety level 4 (BSL4) laboratory at National Emerging Infectious Diseases Laboratories, Boston University (Boston, MA).

### Calculation of infection rate

The rate of infected cells was calculated as follows. At an appropriate time after infection, cells were fixed with 10% neutral buffered formalin (NBF; LC146705; ThermoFisher Scientific) overnight at 4°C.For experiments using GFP-EBOV, cells were stained with Hoechst 33342 dye for 30 minutes at room temperature. Cells infected with WT-EBOV were fixed in 10% NBF overnight at 4°C, followed by permeabilization in 0.2% TritonX-100 (215682500; ThermoFisher Scientific), and then blocked with 3.5% Bovine Serum Albumin (BSA) (BP1600; ThermoFisher Scientific) for 1 h at room temperature. After washing with PBS twice, the cells were stained with anti-EBOV GP antibody (4F3) diluted 1:1,500 in 3.5% BSA for 1 h. A goat anti-mouse secondary antibody conjugated to Alexa Fluor 488 was diluted 1:2,000 in 3.5% BSA and incubated with cells for an additional 1 h. Nuclei were stained with Hoechst 33342 dye. Plates were imaged using a Cytation 1 multimode plate reader (Biotek, Winooski, VT). Multiple non-overlapping images were taken for each well to ensure at least 5,000 cells were captured. The number of GFP or GP-positive cells and nuclei in each well were counted using CellProfiler (Ver. 4.2.1; Broad Institute, Cambridge, MA) pipelines developed by the authors (available upon request). The infection rate, defined here as the percentage of GFP- or GP-positive cells per well, was determined by dividing the number of positive cells by the nuclei count. Reported means and standard deviations are for three biological replicates.

The yield assay of released viral particles was performed using Vero E6 cells incubated with 8-fold serial dilutions of the supernatantfrom samples at 37°C for 48 h. Virus release efficiency, defined as the number of GP-positive cells per nuclei, was calculated as described above., Results are reported as the rate of infection per mL of inoculated supernatant.

### Small interfering RNA (siRNA)

siRNAs targeting several exons of CAPG were purchased from Qiagen (Germantown, MD) (**Supplementary Table 1**). AllStars Negative Control siRNA (nonsilencing siRNA; 1027281) was purchased from Qiagen and used as a negative control. HeLa cells were transfected with respective siRNAs at several concentrations (see figure legends) using lipofectamine RNAiMAX (13778075; ThermoFisher Scientific) following the manufacturer’s protocol. After a 48h incubation the cells were either infected in the BSL4 laboratory or further transfected with DNA plasmids. ..

### Generation of knockout (KO) clonal cell lines

Lentiviral vectors encoding CAPG sgRNAs (clone ID HS5000023627 GCTCATCCCGGGATGACTGCTGG; clone ID HS5000023628 GTTGAGGTGCACAGCCAGCACGG Sigma-Aldrich) were transduced into HeLa cells at MOI of 1.0 with 8 μg/ml of polybrene (Sigma-Aldrich) by centrifuging cells and vectors at 2,300 x rpm for 1h at 22°C. After further incubation with 5 μg/ml puromycin (A11138-03; Gibco) for 2 h at 37°C, cells were washed with PBS, then incubated in fresh DMEM+10% FBS. At 48 h post transduction, cells were trypsinized and seeded into 96 well plates (1.0×104 cells/well) in DMEM+10% FBS supplemented with 20 μg/ml G418 to select for transduced cells.Selection was repeated for 6 passages and selected cells were expanded into 24 well plates and transfected with pX330-U6-Chimeric_BB-CBh-hSpCas9 (#42230; Addgene, Watertown, MA) by Trans IT LT-1 (Mirus Bio, Madison, WI). Single cell clones were established by the limit dilution method. Knockout was confirmed by both Sanger sequence (conducted by GENEWIZ, NJ, USA) and immunoblotting.

### Cell viability assay

The effect of CAPG depletion on viability of KO and siRNA-treated cells was assessed at several timepoints using a 3-[4,5-dimethylthiazol-2-yl]-2,5 diphenyl tetrazolium bromide (MTT) assay (11465007001; Sigma-Aldrich) per manufacturer directions. Briefly, 10 μl of MTT labeling reagent was added to each well and incubated for 4 h at 37 °C. 100 μl of solubilization buffer was added and incubated for additional 1 h. The spectrophotometrical absorbance was measured by Tecan Spark plate reader V2.1 (Tecan, Switzerland) at 575 nm wavelength with a reference wavelength of 675 nm. Cell viability for each well were normalized by subtracting the reference measurement from the 575 nm absorbance value.

### VLP Release Assay

HeLa cells were seeded in 6 well plates (2.0×10^5^ per well) and incubated overnight before transfection with pCAGGS-VP40 using lipofectamin LT-1 (MIR2304; Mirus Bio) according to the manufacturer’s protocol. Cell lysates and supernatants were harvested48 h after transfection. After 48h, 2 mL supernatant samples from each well were harvested and clarified by low-speed centrifugation (3,000 x g, 15min).The clarified supernatant further concentrated and purified over a 20% sucrose cushion in a 60 mm polypropylene tube (328874; Beckman Coulter, Brea, CA), which was spun in a ultracentrifuge at 30,000 x g for 2h at 4°C (SW 60 Ti rotor, Beckman) The pellet was resuspended in 20 μl of PBS for 1 h at 4 °C, and viral protein content was assessed by immunoblot. The remaining cells were lysed using RIPA buffer and used in immunoblot assays.

### Immunoblot assay

Cell lysates were prepared as follows: After removing medium and washing with PBS twice, an appropriate volume of RIPA buffer (BP-115; Boston BioProducts, Milford, MA) with protease inhibitor (A32965; ThermoFisher Scientific) was added to adherent cells. The cells in the buffer were physically disrupted by repeated freeze-thaw cycles and pipetting. The lysates were cleared by centrifugation and supernatants were collected for subsequent immunoblotting. Lysate samples from EBOV-infected cells were removed from the BSL4 following virus inactivation. Briefly, sodium dodecyl sulfate (ThermoFisher Scientific) was added to cell lysate for a final concentration of 1%, and this mixture was boiled at 100°C for 10 min. Protein content of lysate samples was assessed using a Jess system (ProteinSimple, San Jose, CA) according to the manufacturer’s recommendations. Band intensities were calculated using ImageJ (Ver. 2.3.0).

### Immunofluorescence based detection of virus infection and uptake of virus into cells

For immunofluorescent assays, cells were fixed with 10% NBF overnight at 4°C, then permeabilized with 0.2% TritonX-100 for 15 minutes. After blocking with 3.5% BSA, cells were stained with respective primary antibodies for 1 h at room temperature.Following PBS washes, secondary antibodies were added for an additional 1 h. The cells were further stained with HCS CellMask Blue (H32370; ThermoFisher Scientific) to visualize both cell bodies and nuclei. After washing with PBS thoroughly, the cells were imaged using a Ti2 Eclipse microscope (Nikon, Tokyo, Japan) with a100x oil immersion lens.

Viral internalization assays were conducted according to the method described in [60,63]. Briefly, HeLa cells were seeded in 8-well chamber slides (80826; Ibidi, Gräfelfing, German) at 2.0×10^4^ cells per well and incubated with WT-EBOV for 1 h at 14 °C to synchronize the timing of binding of viral particles to the cell surface. Immediately after the incubation, cells were washed three times with cold PBS, then incubated in fresh DMEM with 10% FBS for 6 h at 37°C. After fixation with 10% formalin, the cells were blocked in 3.5% BSA and stained with the anti-GP antibody (4F3) for 1 h without permeabilization. A goat-anti mouse secondary antibody conjugated to Alexa Fluor 594 was used to label membrane-bound GP. Then cells were permeabilized, blocked again, and restrained with the same anti-GP antibody, followed by incubation with Alexa Fluor 488-conjugated goat anti-mouse IgG antibody. The cell body and the nuclei were stained with HCS CellMask blue. Images were taken at multiple z-planes on a Nikon Ti2 and deconvolved using Microvolution software (Microvolution Inc., Cupertino, CA) run on Fiji [64]. The deconvolved images were analyzed using Imaris 3D image analysis software (Bitplane Inc., Belfast, UK). Green foci and merged green-red foci were counted and virus internalization efficiency was calculated as the ratio of green foci to total foci.

### Proximity ligation assay

The Duolink Proximity Ligation Assay kit (Sigma-Aldrich) was used to observe proximity between viral proteins inside cells In brief, HeLa cells were seeded onto 18 well chamber slides (81816; ibidi) at 0.5×10^4^ cells per well. After incubation overnight, the cells were infected with WT-EBOV for 24 h. The cells were fixed and permeabilized, and the proximity ligation was performed following the manufacturer’s protocol using specific primary antibodies described above. Actin filaments were stained with Alexa Flour 488-conjugated phalloidin (A12379; ThermoFisher Scientific) for 30 min. Red foci inside cells indicated that two targeted proteins were in close proximity. Images were taken at 40x or 100x with oil immersion on a Nikon Ti2 microscope. 3D images were deconvolved as above and used to detect protein interactions with phalloidin-labeled actin filaments.

### Tripartite split-GFP assay

HeLa cells were transfected with GFP B10, GFP B11 and GFP1-9-T2A-mCherry expression plasmids at a ratio of 1:5:5 (typically 15 μg B10 plasmid, 75 μg B11 plasmid, and 75 μg pCDNA3-GFP1-9 T2A mCherry per transfection) using Lipofectamine 2000 (Invitrogen) according to the manufacturer’s instructions. Four transfections were performed for each combination. Cells were incubated for 48 h, fixed with 4 % paraformaldehyde, stained with Hoechst 33342 dye and stored in the dark at 4° C until imaging. Fixed cells were imaged on a PerkinElmer Opera Phenix High Content Screening System using a 10x/air NA 0.3 confocal lens and acquiring nine fields per well without stacking. The excitation and emission wavelengths (Exc/Em) for each channel were: Hoechst33342 (375/435-480), EGFP (488/500-550), and mCherry (561/570-630). The EGFP channel was scanned independently whereas mCherry and Hoechst were acquired simultaneously. After applying basic flatfield and brightfield corrections, image analysis was performed using Harmony High-Content Imaging and Analysis Software (version 4.9). First, nuclei were identified by mean intensity with area < 300 μm^2^, intensity < 6000 and roundness < 1.5 using method C. Second, transfected cells expressing mCherry were delineated using mean intensity with area > 100 μm^2^ using method B. Finally, total GFP spots in the transfected cell region of interest were enumerated based on intensity using method D. The nine fields from each well were merged without overlaps and the calculated output values were exported for statistical analysis in GraphPad Prism (GraphPad Software, San Diego, CA).

### Real-Time quantitative PCR (RT-qPCR)

The quantification of viral and cellular RNA from both supernatants and cell lysates was performed as described below. In brief, the cell supernatant was collected and centrifuged to eliminate cell debris. TRIzol LS reagent (10296028; ThermoFisher Scientific) was added to the supernatant at a 3:1 ratio. Cell lysate was collected by lysing cells with TRIzol reagent (15596018; ThermoFisher Scientific) after supernatant had been removed and cells had been washed in PBS. Samples were stored at −80°C until use. Viral and cellular RNA was extracted using phenol-chloroform separation method following the manufacturer’s protocol. RT-qPCR was performed using Luna^®^ Universal Probe One-Step RT-qPCR kit (E3006L; New England Biolabs (NEB), Ipswich, MA) and amplification was detected and validated by CFX96 Touch Real-Time PCR Detection System (Bio-Rad, CA, USA). The cycling protocol was: 55°C for 10 min, 95°C for 1 min, followed by 40 cycles of 95°C for 10 sec and 60 °C for 30 sec. See **Supplementary Table 2** for primer and probe sequence information [65–68]. All the primers and probes were synthesized by Integrated DNA Technologies (Coralville, IA). ΔΔCt was calculated relative to control samples following a calculation of ΔCt from each sample based on the Ct of GAPDH. To generate RNA for a standard curve, 612 bp of NP sequence including the RT-qPCR target region described in Supplementary Table 2was amplified from a pC-NP plasmid using a forward primer containing a T7 promoter (5’-TAATACGACTCACTATAGGGTCTGTCCGTTCAACAGGG-3’) and reverse primer (5’-ATCACAGCATCGTTGGCATCATG-3’)). After electrophoresis and gel extraction by Monarch DNA Gel Extraction Kit (T1020S; NEB), the PCR product was transcribed using HiScribe T7 High Yield RNA Synthesis Kit (E2040S; NEB) according to the manufacturer’s instructions, followed by degradation of the DNA template by DNaseI treatment (M0303S; NEB) and purification using Monarch Cleanup Kit (T2040S; NEB). The approximate RNA copy number was calculated based on its molecular weight and absorbance measured by a NanoDrop 1000 (ThermoFisher Scientific). For each RT-qPCR reaction set, a five-point 10-fold diluted standard was included to ensure performance of the assay.

### Statistical analysis

GraphPad Prism version 9.0.0 for Mac (GraphPad Software, San Diego, CA) was used to carry out one-way ANOVAs with Dunnett’s or Tukey’s multiple comparisons test, and p-values from these analyses were used to determine statistical significance. Significance was taken as P <0.05.

## Results

### Suppression of CAPG expression impairs EBOV infection

The importance of CAPG for EBOV infection was initially tested by suppressing its expression using four siRNA targeting different portions of the CAPG mRNA. Each reduced CAPG expression ranging from 60 to 80% loss (**Fig. 1A**). Virus yield was measured by challenging cells at an MOI of 0.01, sampling medium after 48 h and measuring virus titer on fresh Vero cells. Compared to non-targeting siRNA, virus yield was greatly reduced (>90%) for each CAPG specific siRNA (**Fig. 1B**). The impact on infection efficiency was confirmed using recombinant EBOV, encoding GFP as an infection marker, and cells fixed after 48 h, which after accounting for maturation of GFP, corresponds to approximately 2 rounds of virus replication [69]. In each case a significant reduction (P<0.001) of EBOV infection ranging from 60 to 80% was seen (**Fig. 1C, D**) but was not as great as seen for the virus yield experiment. Cell viability was confirmed at several time points after transfection of siRNA, and no significant difference was observed compared to non-targeting siRNA. (**Fig. S1B**). Taken together, our initial data indicated a role for CAPG in controlling early infection as well as egress but having a much greater impact on virus yield from infected from cells.

**Fig. 1.**
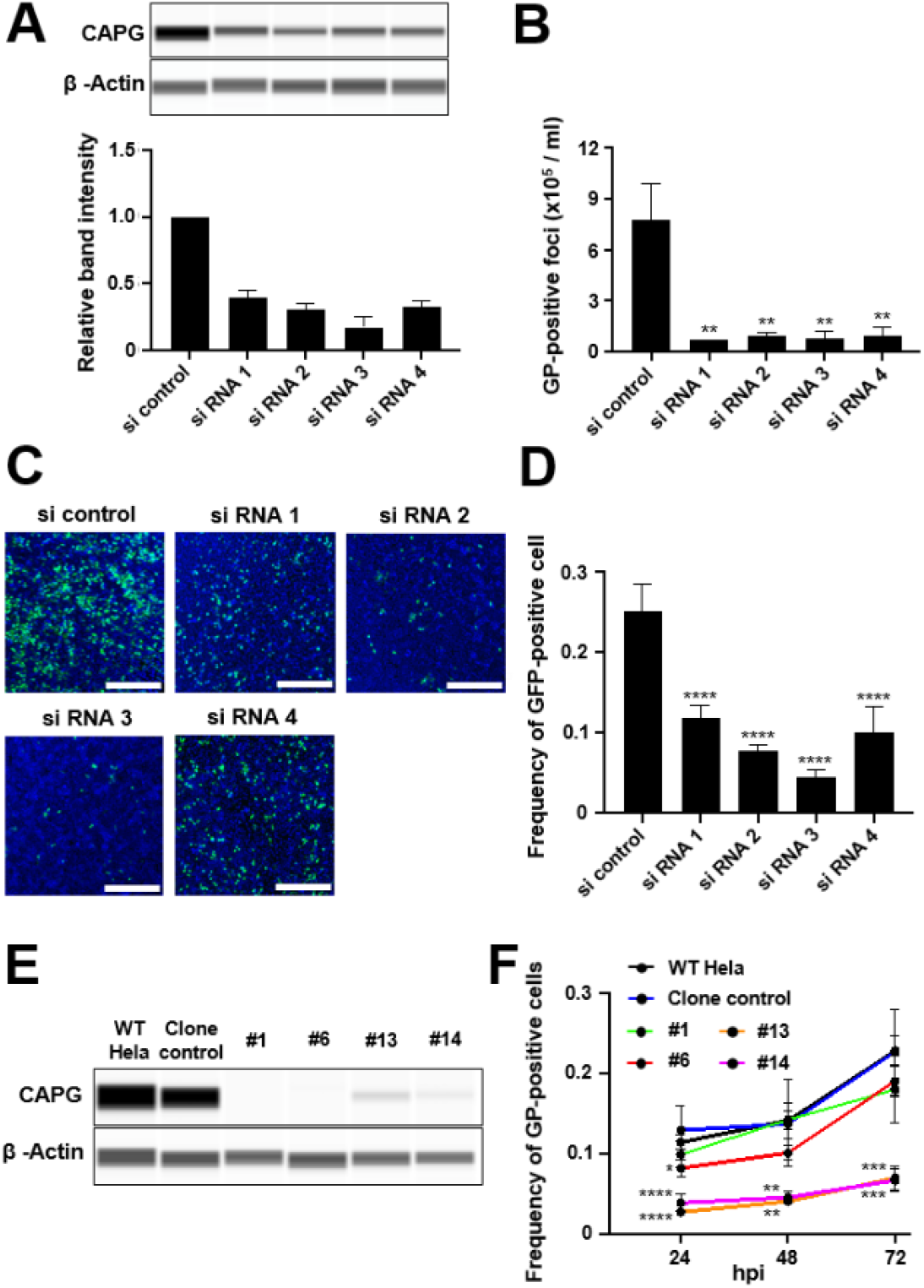
Suppression of CAPG impairs EBOV infection. **A.** CAPG knockdown was confirmed by immunoblot assay. Four siRNA targeting different regions of Human CAPG were transfected into Hela cells at 40 nM. After 48 h cells were challenged with virus and a replicate set were lysed and protein expression analyzed by immunoblot. ß-actin was used as a loading control. Band intensity from triplicate samples was calculated using ImageJ, and normalized to the β-actin loading control band intensity. **B.** Yield of infectious particles in the supernatant was measured by counting GP positive foci on VeroE6 cells. Cells were inoculated with the supernatant from wild-type EBOV-infected cells by limiting dilution. At 48 h after inoculation, cells were fixed and stained with anti-GP antibody and foci counted. **C.** siRNA treated cells were challenged with GFP-EBOV with representative images shown from 48 hpi. Infected cells expressing GFP (green) and nuclei stained with Hoechst 33342 (blue) are visible. Scale bars =500 μm. **D.** Count of GFP-positive cells. The numbers of GFP-positive cells and the nuclei in each image were counted by CellProfiler software and infection efficiency was calculated by dividing the number of GFP-positive cells by that of the nuclei. **E.** CAPG expression in each KO and KD clones was detected by immunoblot. β-actin was used as a loading control. WT Hela = parental WT Hela cells and clones are indicated. **F.** Time-course analysis of virus spread each clone. At each indicated time point after infection with WT-EBOV, the cells were fixed and stained with anti-GP antibody. The number of GP-positive cells were counted by CellProfiler software, then normalized to that of the nuclei count in the same image. All data are representative of three independent experiments. One-way ANOVA with Dunnett’s multiple comparisons test was used for statistical analysis relative to non-targeting siRNA treated control samples or the CRISPR clone control. **, P<0.01; ***, P<0.001; ****, P<0.0001.

To validate the siRNA findings, CRISPR/Cas9 was used to establish two knockout (KO) cell clones (labeled as #1 and #6), and two partial CAPG-expressing clones (>90% loss of CAPG, #13 and #14) that mimicked the incomplete siRNA induced suppression of CAPG. Each clone had alterations in exon 4, which encodes amino acid residues 66 to 117 and the first actin binding domain (see NCBI NM_001747.4 and ref. [70]). Sequencing indicated clones #1 and 6 had disruption of all alleles while clone #13 and #14 maintained at least one functional allele (**Fig. S2**) and was reflected in loss or reduced protein levels on immunoblots (**Fig. 1E**). Controls were parental Hela cells (WT Hela) and a control line that was CRISPR/Cas9 treated but had no change in CAPG expression. Virus replication was assessed over different times (up to 72 h) by measuring spread of virus through the cell culture by staining for expression of the EBOV glycoprotein (GP). The two clones that had very low but detectable expression of CAPG showed a 4-fold block in virus infection, with replication remaining low at all time points (**Fig. 1F**). Interestingly, the two clones that had a loss of detectable CAPG expression generally showed normal infection levels with a difference in infectivity that was significant only at the 24 h time point for clone #6, suggesting these had compensated for loss of CAPG expression which is known to occur during KO selection process for some other cell genes [71]. The growth of each clone was similar to wild type cells (**Fig. S1C**). Furthermore, phalloidin staining patterns, measuring F-actin content and morphology in cells appeared similar to wild type cells (**Fig. S3**)suggesting that loss of CAPG did not have global effects on F-actin. Overall, our data indicates that suppression of CAPG by either CRISPR or siRNA results in loss of infectivity, with an apparent greater effect on virus spread in infected cell cultures.

### CAPG suppression does not affect virus entry and early mRNA production in cells

To examine early defects in the virus infection cycle, virion binding and uptake into cells was measured. Virus uptake into cells was measured 6 h after initiation by staining fixed cells with a sensitive GP-specific antibody followed by different fluorescently colored secondary antibodies before (red) and after (green) permeabilization of the plasma membrane with a non-ionic detergent. The different colored secondary antibodies allowed differentiation of surface bound (both red and green fluorescent virions) versus those that had entered cells and were inaccessible to labeling until membrane permeabilization (green only). Despite a reduction in the internalized virions for siRNA 1 and 2, this change was not significant (**Fig. 2A**). As a second measure of virus uptake, siRNA treated cells were challenged with WT-EBOV and RNA extracted at 4 h post-infection, a time soon after entry into the cell cytoplasm and at the earliest time when virus mRNA can be detected [69]. EIPA and CPZ were used as active and inactive control inhibitors for macropinocytosis (major entry path used by EBOV) and clathrin-dependent endocytosis (not substantially used by EBOV), respectively [72,73], being treated 1 h before incubation with virus. RT-qPCR was performed targeting the NP gene. While EIPA treatment significantly reduced viral RNA levels (P<0.01), each siRNA treatment cells showed no significant difference to either non-targeting siRNA treated or chlorpromazine treated cells at this time (**Fig. 2B**). These findings are consistent with CAPG having only a minor role in virus uptake into cells.

**Fig. 2.**
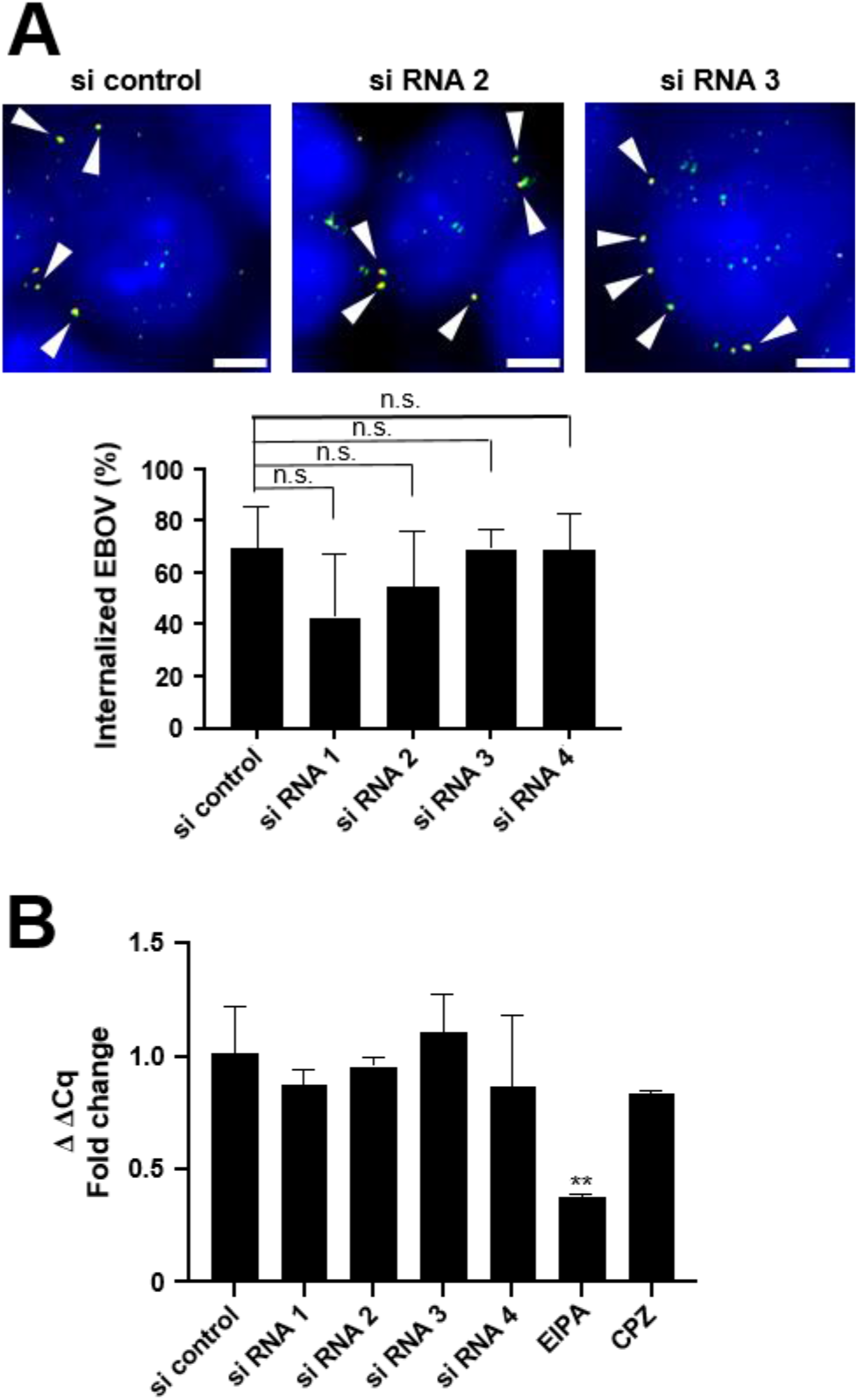
Effect of CAPG suppression on EBOV uptake into cells. **A.** Hela cells were treated with siRNA for 48 h, incubated with WT-EBOV at 14°C for 1h and then 6 h at 37°C to give synchronized virus uptake. Cells were then fixed and stained with anti-GP and Alexa Flour 594 antibody (red). Subsequently, cells were permeabilized with 0.2% TritonX-100 for staining internal viral particles with anti-GP and Alexa Fluor 488 antibody (green). HCS CellMask Blue was used for staining both the cell cytoplasm and nucleus (blue). The percentage of internalized virus was calculated as the ratio of green foci against total foci and shown in the graph below the images. Arrowheads show double stained puncta. Scale bars = 5 μm. **B.** Real-Time quantitative PCR (RT-qPCR) analysis to detect viral RNA early during infection. siRNA-treated Hela cells were incubated with WT-EBOV for 1 h, then the cells were washed and fresh medium was added onto the cells. The cells were lysed at 4 hpi for RNA extraction. Viral RNA was detected using a primer and probe set targeting NP gene (see Supplementary Table 2). ΔΔCt was calculated using GAPDH as a reference control in each sample. The data are shown as fold change relative to siRNA non-targeting control. A 1 h pre-treatment with 50 μM of Ethylisopropylamiloride (EIPA) or Chlorpromazine (CPZ) were used as macropinocytosis and clathrin-dependent endocytosis inhibitors, respectively.

### CAPG is required for efficient production of virus from infected cells

Since actin function is known to be important for EBOV budding from cells [30,45,46] and CAPG regulates actin polymerization, we next tested whether CAPG depletion impacted budding of virions from cells. Viral RNA levels present in cell culture medium was used to gage virion release from cells. Signals were compared between viral RNA in supernatants and cell lysates (**Fig. 3A**). While this measurement was complicated by the previously measured reduction in initial infection efficiency, in each case, levels of virus RNA in supernatants were reduced in supernatants (increase in ΔCq) relative to cell lysate levels and was more pronounced for the weaker siRNA #1 followed by #3 and #4.

**Fig. 3.**
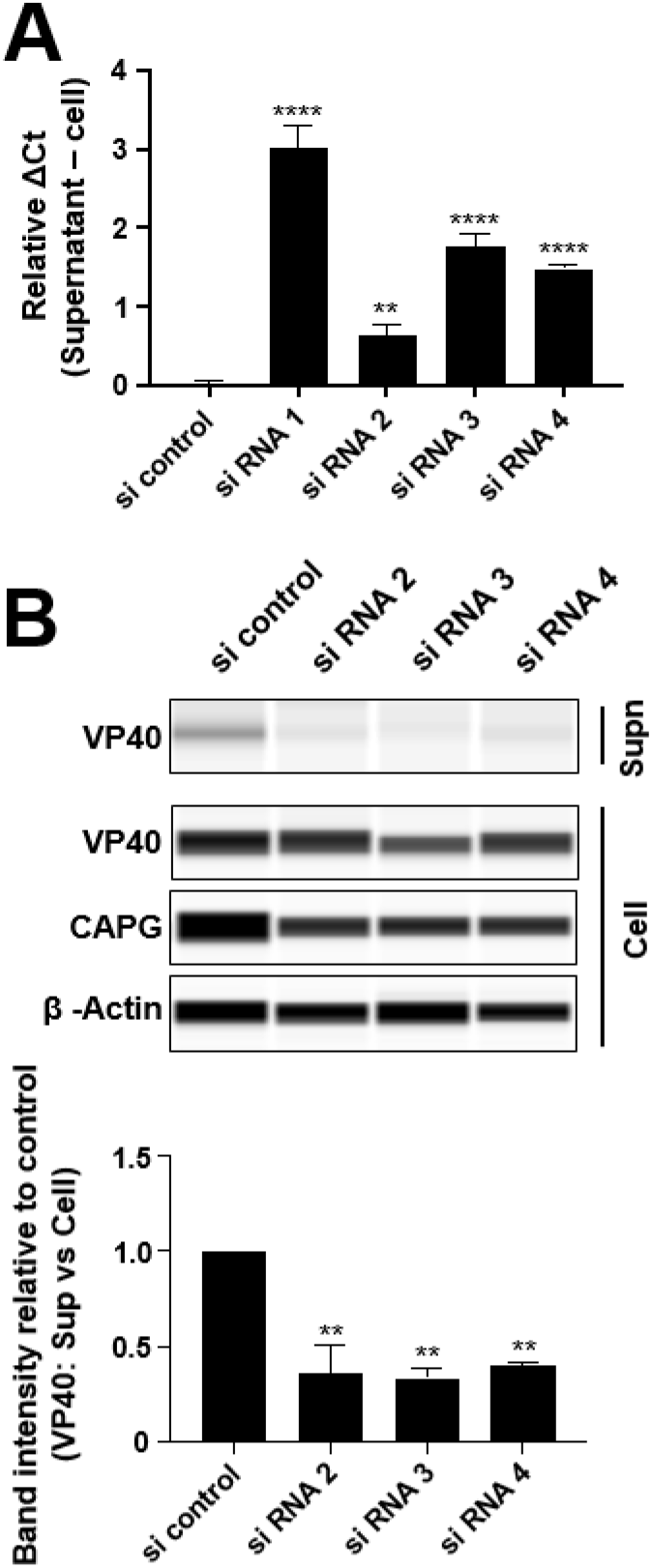
Effect of CAPG suppression on EBOV release from cells. **A.** Measurement of the quantity of viral RNA released from siRNA treated cells. At 48 hpi, RNA was extracted from the supernatant and the remaining cells, then virus RNA levels measured by RT-qPCR using primers for NP. The graph indicates ΔCt (supernatant – cell) in each sample. **B.** The efficiency of VLP formation from cells treated with each indicated siRNA. Hela cells seeded in 6 well plate were transduced with siRNA (40 nM each) and pCAGGS-Ebola VP40 plasmid (0.5 μg) using lipofectamine. At 48 h post transfection, the supernatant was collected and centrifuged to remove cell debris. VLPs were collected by pelleting through a 20% sucrose cushion. VLP pellets and cell lysates were analyzed by immunoblot. Band intensity from each sample is shown relative to siRNA non-targeting control. All assays were repeated at least twice and the representative data sets were shown in this figure. One-way ANOVA with Dunnett’s multiple comparisons test was used for statistical analysis relative to control samples. The means of two or three independent experiments ± SDs is shown. n.s., non-significant; **, P<0.01; ****, P<0.0001.

Since Ebola virus-like particles (VLPs) can assemble and bud from cells through expression of EBOV VP40 alone, Hela cells were transfected with siRNA and VLPs were recovered from culture supernatants by pelleting through a sucrose cushion. VP40 levels in both cell lysate and collected pellets were measured by immunoblot (**Fig. 3B**). Compared to VP40 expression in cell lysates, VP40 in supernatants was significantly reduced (P<0.01) by 65-70% in cells treated with each of the strongest acting siRNA (#2-4). This indicates that VP40 release from cells and therefore budding of virus from cells is abrogated upon CAPG depletion. In summary, budding of EBOV becomes inefficient in CAPG deficient cells and is reflected in VP40 release as VLPs.

### VP40 and GP are found in close proximity to CAPG in the cytoplasm

Since loss of CAPG was sufficient to reduce release of VP40 from cells, we tested if CAPG and VP40 associate in cells. To analyze the physical interaction between viral proteins and CAPG, we performed proximity ligation assays which are reported to give signal when two targeted proteins are within 40 nm of each other [74] using wild type EBOV-infected Hela cells. As a positive control, we checked amplified signals from NP and VP30 which are known to interact [75,76]. As shown in **Fig. 4A**, amplified signals (red) were clearly observed when probed with specific antibodies for each native protein. Similarly, for VP40 and CAPG, strong signals were seen within the cell cytoplasm. To gage when this interaction took place, we tested GP association, a protein that meets with maturing virus particles late in virus assembly. Again, distinct signal was observed that extended to the periphery of cells. However, little signal was seen when VP35 was targeted together with CAPG, suggesting that CAPG does not closely associate with VP35, which becomes part of the nucleocapsid at early steps of virus assembly. These results indicate that GP and VP40 are present at sites that are in close proximity to CAPG. One explanation was that CAPG and VP40 co-associate through actin. Indeed, samples stained with phalloidin to detect F-actin showed localization of the CAPG-VP40 puncta along actin filaments and in cell projections, that may represent filopodia (**Fig. 4B**) and known to be sites of EBOV budding [32,77]. These results indicate that CAPG localizes with VP40, potentially along actin filaments and is also associated with GP, suggesting a relationship to late stage maturing virions.

**Fig. 4.**
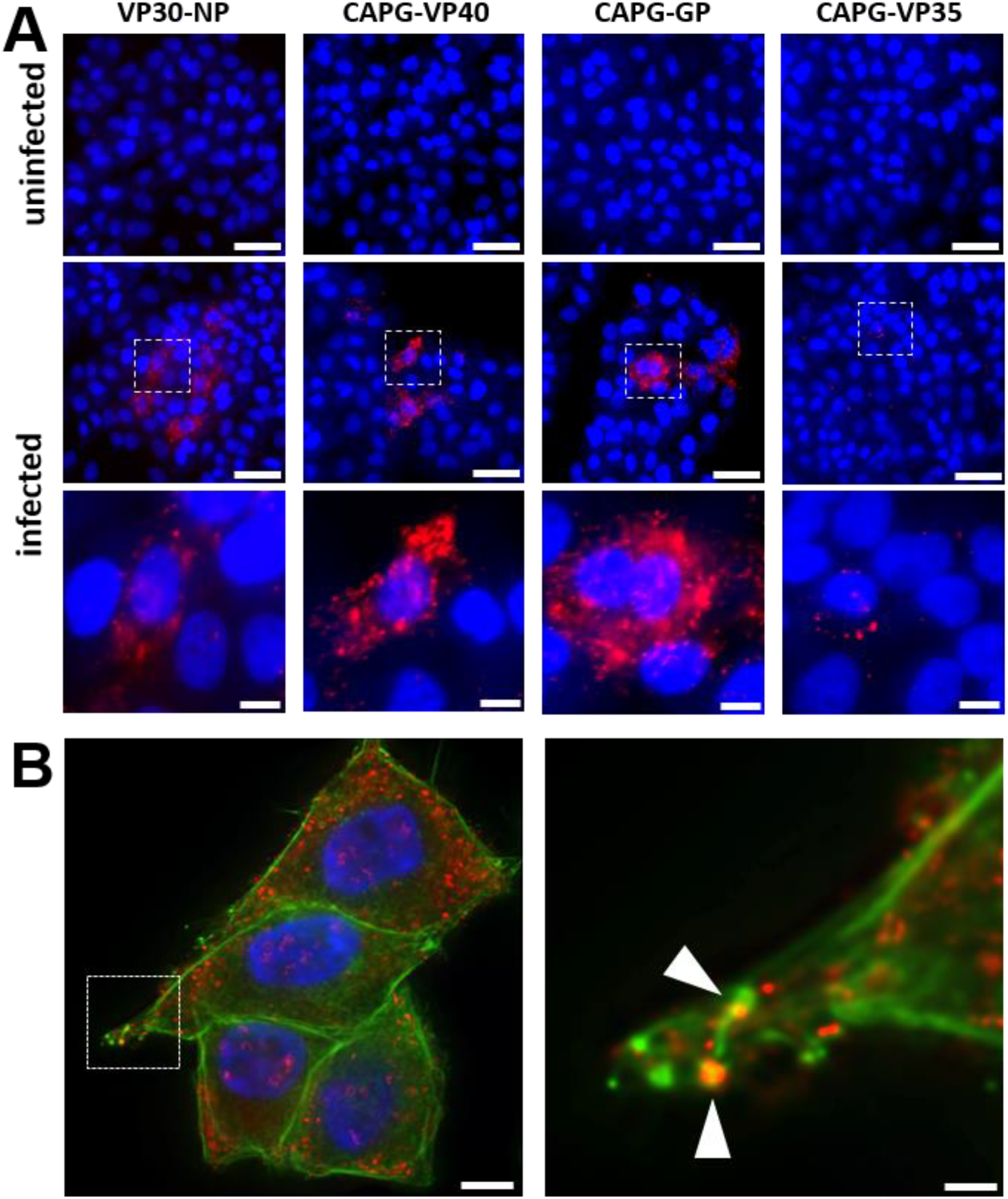
EBOV proteins localize in close proximity to CAPG in the cytoplasm. **A.** Proximity ligation assay was performed to detect CAPG and viral protein association. Hela cells were infected with WT-EBOV for 24 h, then fixed and permeabilized. Cells treated with the indicated antibodies specific for each native protein and detected protein complexes stained red. Cell nuclei were stained with Hoechst 33342 (blue). The lower set of images from infected cells are magnified from the indicated (dotted squares) sections of images. Scale bars = 50 μm, and 10 μm in magnified images. (**B**)A representative image (left) of amplified signal (red) of VP40-CAPG and phalloidin (green). The images were the mid set of images from a z-stack. A magnified image (right) is shown for a region indicated by the dotted square. Arrowheads indicate sites where CAPG-VP40 complexes and and phalloidin staining (F-actin) overlap. Scale bar = 10 μm, and 2 μm in magnified image.

### Identification of CAPG subdomains involved in actin interaction are important for association with VP40

CAPG has a unique feature among the Gelsolin/villin family proteins consisting of three main structural repeats of single gelsolin-like domains (**Fig. 5A**) S1, S2, and S3 [78]. To identify subdomains in CAPG important for association with VP40, we used a split GFP system (**Fig. 5B**) where simultaneous close tethering of three GFP subdomains results in GFP reconstitution and fluorescence [79]. VP40 was tagged with GFP10, and host proteins, CAPG, TSG101, UtrCH and PABPC1 were tagged with GFP11. PABPC1 and TSG101 were used as negative and positive controls respectively as TSG101 is known to associate with VP40 [80] and PABPC1, a polyA binding protein, is not reported to associate with VP40. Each was transfected into cells along with plasmid encoding the third portion of GFP, GFP1-9 that was expressed together with an autocleaved mCherry protein that served as a transfection efficiency control. GFP11 tagging was performed at either the N or C-terminus. The number of green (containing reconstituted GFP) fluorescent cells was normalized to the number of red fluorescent (mCherry expressing) cells.

**Fig. 5.**
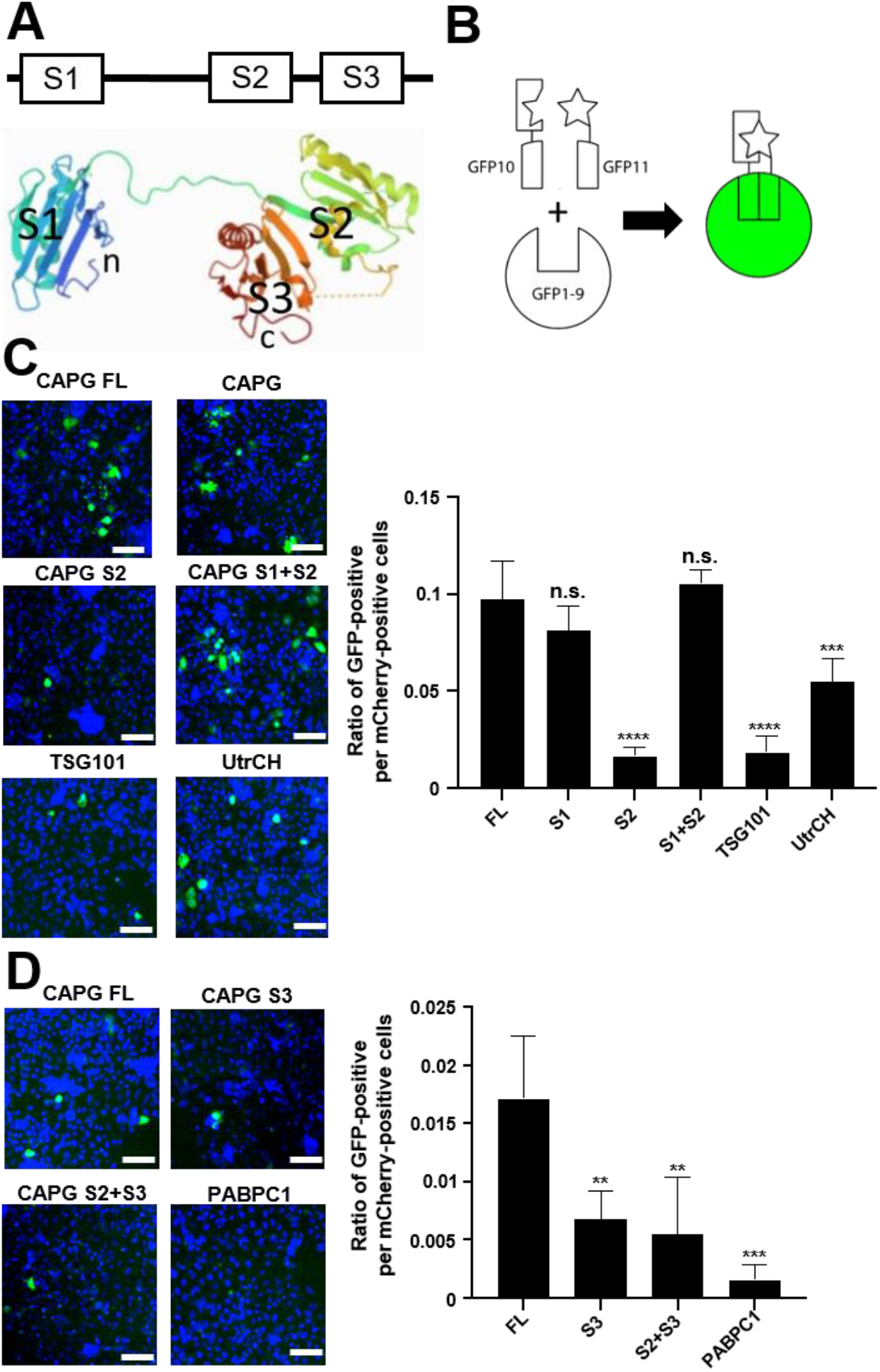
Identification of CAPG subdomains important for interaction with VP40. **A.** Schematic representation of the CAPG protein. Each gelsolin domain is indicated by S1, S2 and S3. Lower panel shows the 3D structure of CAPG derived from PDB 1JHW. **B.** Shows the arrangement of the tripartite split GFP system. For this assay, the GFP10 strand is used to tag VP40 and GFP11 strand was used for the indicated host cell protein targets. When each part is brought together with the remainder of GFP, cells fluoresce green through formation of GFP. **C and D.** At left are representative images showing cell nuclei stained with Hoechst 33342 (blue) and GFP positive cells (green). At right is the quantitation of the images. **C.** are N-terminal fusions with GFP11 peptide and **D.** are C-terminal fusions. The ratio of GFP-positive per mCherry-positive cells is shown. GFP-positive cells and mCherry-positive cells were counted by CellProfiler software, then the number of GFP positive cells were normalized that of mCherry-positive cells in each image. The means of four independently treated samples ± SDs are shown. **, P<0.01; ***, P<0.001; ****, P<0.0001 by One-way ANOVA with Dunnet’s multiple comparisons test.

When full-length CAPG was used as bait, approximately 10% of cells expressing mCherry also showed strong green fluorescence and was similar to that seen with the S1 domain alone or the S1 and S2 domains together (**Fig. 5C**). In contrast, the S2 domain alone gave little green fluorescence. TSG101, also gave low fluorescence but was significantly above the background of the assay. The S3 domain alone was not well expressed when N-terminal tagged but did tolerate C-terminal tagging. Despite lower overall signals for the C-terminal tags, both S3 alone and S2 together with S3 gave little signal that was 3-fold less (P<0.01) than the full length construct (**Fig. 5D**). The c-terminal domain of utrophin (UtrCH) was used to determine signal for a known actin binding protein. UtrCH binds actin monomers within F-actin in a ratio of 1:1 and tagging with GFP does not disrupt this interaction [81]. While, signal was present, it was half that seen and significantly less (P<0.01) than that from full length or the CAPG S1 domain alone (**Fig. 5D**) suggesting the CAPG-VP40 was stronger. These results indicate that the S1 region, which contains one of two actin binding motifs of CAPG [82,83], is required for association with VP40.

## Discussion

The Ebola virus replication cycle is heavily dependent upon actin [25]. Early work with inhibitors of actin polymerization such as cytochalasin D or latrunculin, showed dependence for entry by micropinocytosis as well as for virus assembly [30,84]. Similarly, actin-regulating host proteins such as RhoA and the ARP2/3 actin nucleating complex have been identified as necessary for these steps [28,29]. In addition, actin filament plays an important role in transporting the replication complex and interactions with VP40 in the maturing virion [32,45,47]. Unfortunately, the heavy involvement of actin in many cellular processes, makes interpretation of findings from direct manipulation of actin highly complicated. However, recent use of actin-accessory proteins, such as those that bind and stabilize F-actin, has allowed advances in understanding actin function and dynamics [81]. In this study, we studied the importance of the actin-accessary protein, CAPG, to gain insight into the role of actin in EBOV infection. Unlike other accessory proteins, like utrophin, that bind monomers along the actin filament, CAPG is present only on the growing end of F-actin and so may offer new insights into how this aspect of actin polymerization impacts virus replication.

Our knockdown and knockout experiments showed the importance of CAPG on infection for both virus entry and egress. In our work, knockdown by siRNA or CRISPR was effective to suppress viral infectivity. Despite what we and others have shown for EBOV uptake dependence on actin function, through macropinocytosis [24,25], we saw only weak impact on this step with particle uptake into cells not significantly affected. Instead, we saw a heavy dependence for virus egress, consistent with previous reports reporting the importance of actin functionality for this step [30,33,45–47]. Consistent with this previous work, we saw similar impact on wild type virus as well as VP40 based VLPs. The site of interaction with virus appears to be late in virus maturation as colocalization using the proximity ligation assay showed distinct puncta aligned along actin filaments toward the cell periphery and were also associated with the EBOV GP, a protein that associates with VP40 when the latter interacts with cell membranes during budding. To more precisely determine the exact site of these interactions further study will be needed using techniques such as high resolution microscopy as has been recently reported [85].

In identifying the portion of CAPG important for VP40 association, we observed very strong signals in the GFP trans-complementation assay. The signals from full length or the S1 domain alone was more frequent than that seen for both TSG101 as well as utrophin. TSG101 is known to strongly bind VP40 through its PPXY motifs and gives strong signals in a similar transcomplementation assay [86]. Furthermore, utrophin is found on F-actin in a 1:1 ratio with each actin monomer. While we do not know how well the recombinant utrophin binds to actin, other reports show that it can be used to label the entire length of the actin filament. Together with the proximity ligation assay outcome, our work supports a relatively strong association of VP40 with CAPG and therefore a potential localization of virus at the growing end of actin filaments. Mechanistically, this interaction may enhance the viruses ability to move through and leave cells. Based on our data, we propose that CAPG connects VP40 and the growing actin filament during the egress step of viral particles.

While additional work is needed to fully understand the CAPG-VP40 interaction and its impact on virus egress, we expect our findings will aid in treatment of EBOV disease. CAPG consists of about 1% of total protein in macrophages, cells that are primary targets of infection and effectors of EBOV disease [87–89]. Therefore, suppressing CAPG function could have significant effects on viral dissemination in patients. Recently, a ROCK inhibitor, was shown to indirectly decrease CAPG expression in fibroblast cells and resulted in improved wound healing in heart disease models [90]. It will be interesting to determine if a similar approach may provide improved outcome in a virus disease model.

## Supporting information

Supplemental Table 3. Cloning oligonucleotide sequences

Supplemental Table 4. Plasmid maps

## Author Contributions

Conceptualization, H.M., and R.A.D.; methodology, H.M., J.P.C.,D.L., D.J.L. and R.A.D.; assay development, H.M., J.P.C., C.J.D., R.M.B., J.J.P.; data analysis, H.M. and R.A.D.; data curation, H.M.; statistical analysis, H.M. and R.A.D.; writing H.M., C.J.D, R.M.B, D.J.L and RAD. This work and personnel were supported in part by NIH P01AI120943. All authors have read and agreed to the published version of the manuscript.

## Acknowledgments

We acknowledge the Chemical Genomics Facility at Purdue Institute for Drug Discovery for the assistance in the image acquisition and analysis.

