## Supplemental figures and table 1 and 2 for "CAPG is required for Ebola virus infection by controlling virus egress from infected cells"

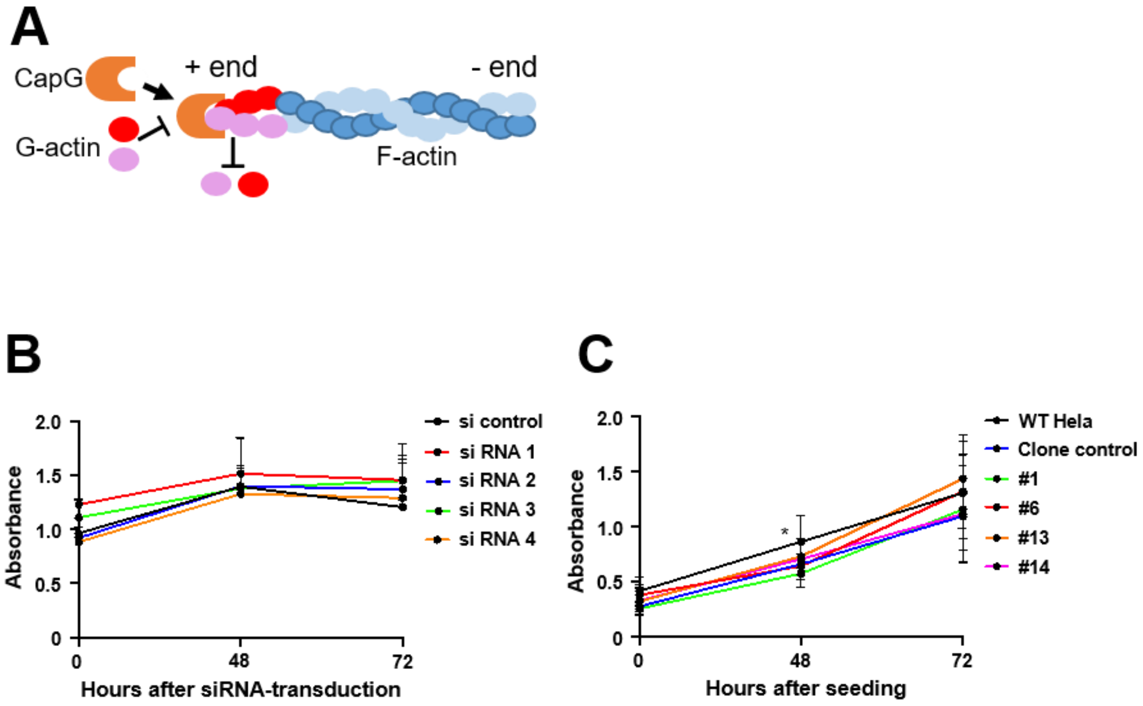


**Supplementary Figure 1. Schematic of CAPG function and cell viability after transduction of siRNA and CAPG knockout cells.** A. CAPG functions to cap the barbed, growing end of actin filaments. This prevents addition of new G-actin monomers as well as their release from already formed filaments. Cell viability of Hela cells after transduction of siRNA **(B)** and knockout (KO) clones **(C)** in a time-course of post-transfection or seeding, respectively. 3-[4,5-dimethylthiazol-2-yl]-2,5 diphenyl tetrazolium bromide (MTT) assay was conducted to check spectrophotometrical absorbance in each sample seeded on 96 well plate. The absorbance was measured at 575 nm, and at 675 nm wavelengths for background detection. Y-axis indicates the absorbance after deduction of measurement at 575 nm wavelength from that at 675 nm. Statistical difference was calculated by One-way ANOVA with Tukey's multiple comparisons test at each time point. *, $p < 0.05$. hpi = hours post infection.

A

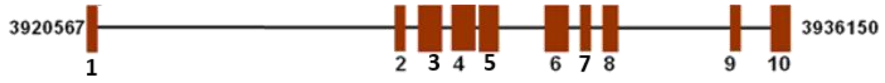


B

#1

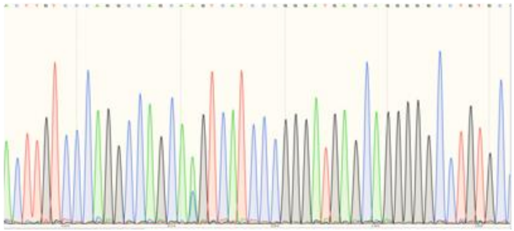

```
nt 243  WT: 5'-gccagcagtcatcccgggatgagcaggggcctgtg
        #1: 5'-gccaAgcagtcatcccgggatgagcaggggcctgtg
        #1: 5'-gccaCgcagtcatcccgggatgagcaggggcctgtg
```

#6

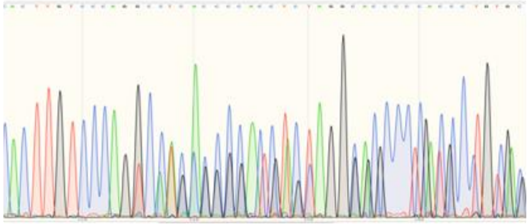

```
nt 243 WT: 5'-gccagcagtcatcccgggatgagcaggggcctgtg
       #6: 5'-gccagc--------cgggatgagcaggggcctgtg
       #6: 5'------cagtcatcccgggatgagcaggggcctgtg
```

#13

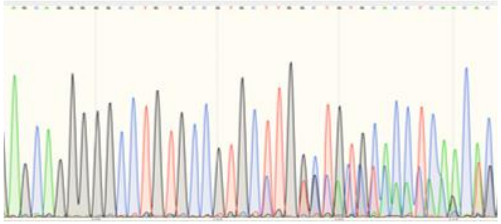

```
nt 272 WT:  5'-gcctgtgccgtgct-ggctgtgcacctcaacacgctg
       #13: 5'-gcctgtgccgtgctTggctgtgcacctcaacacgctg
       #13: 5'-gcctgtgccgtgc----ctgtgcacctcaacacgctg
```

#14

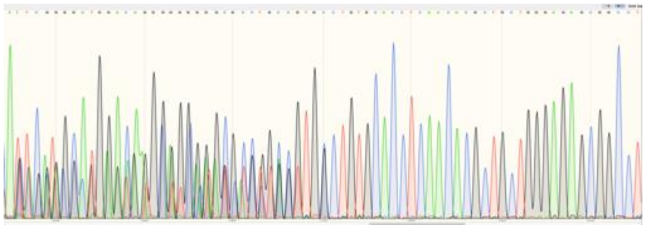

nt 281

**Supplementary Figure 2. Sequence information for INDELs of CAPG KO clones.**

A. Gene structure of CAPG with exons highlighted. Modified from NCBI using CAPG variant 1 NM_0017477.4. B. Sanger sequencing for regions targeted by guide RNA against Exon 4. Below each chromatogram are the sequences inferred for each allele present.

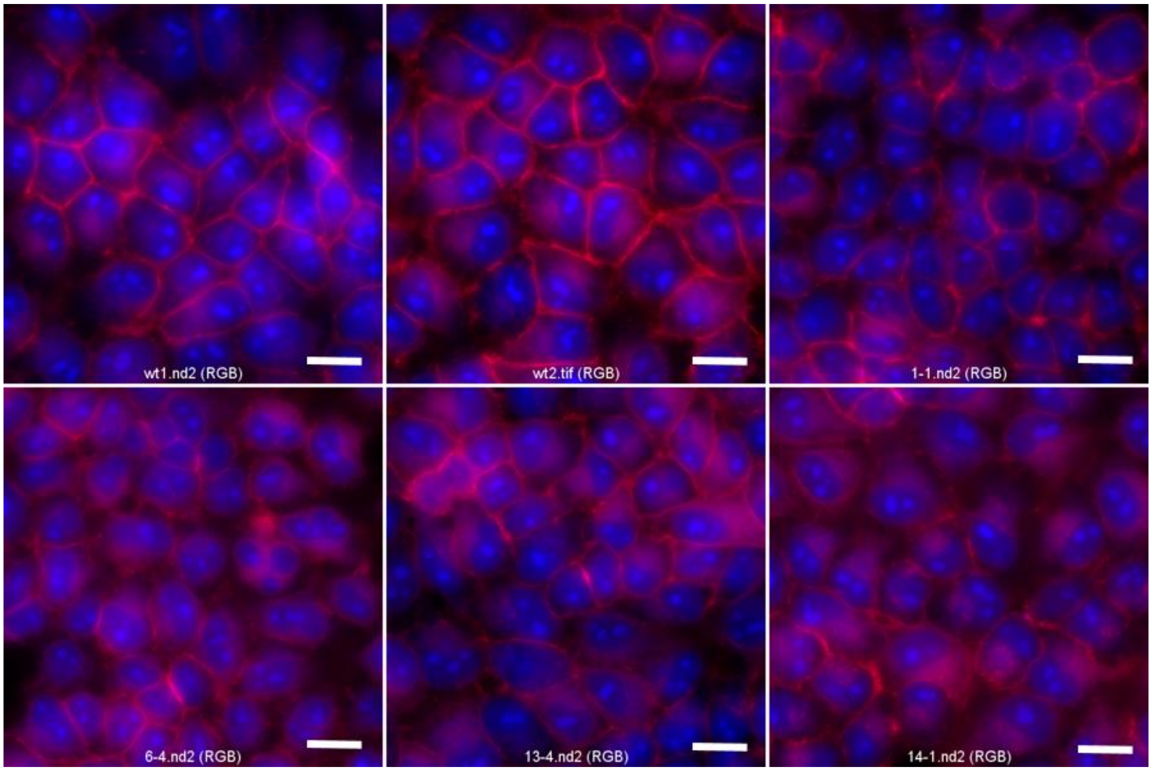


**Supplementary Figure 3. Phalloidin staining of KO and KD cell clones.** Cells were stained with phalloidin to detect F-actin (red) and Hoechst 33342 to detect cell nuclei (blue). Scale bar is 20 µm.

Supplemental tables.

| | Sequence (5' to 3') | Exon |
|---|---|---|
| siRNA 1 | CTGGTGTGGTGGAAAGTCCAA | 6 |
| siRNA 2 | CAGGTGGAGATTGTCACTGAT | 6 |
| siRNA 3 | CGGGCTCTGTGGCAAGATCTA | 8 |
| siRNA 4 | AAGGAGGGCAACCCTGAGGAA | 7 |

**Supplementary Table 1. Sequence information of siRNA used in this study**

siRNAs described here were synthesized by and ordered from Qiagen. siRNA 1 and 4 were customized siRNA, and siRNA 2 and 3 were from FlexiTube siRNA (Cat. No. 1027417). Control siRNA was also a Qiagen product (AllStars Neg. Control siRNA, Cat. No. 1027281). The sequence information of control siRNA is not shown here. The sequence information is shown 5' to 3' (from left to right), and each targeting exons of CAPG gene are also informed here.

| Host | Gene target | Primer/Probe | Amplicon (bp) | Sequence (5'-3') | Reference |
|---|---|---|---|---|---|
| ZEBOV | **NP** | Forward | 80 | GCAGAGCAAGGACTGATACA | **44** |
| | | Reverse | | GTTCGCATCAAACGGAAAAT | |
| | | Probe | | FAM-CAACAGCTT-ZEN-GGCAATCAGTAGGACA-IABkFQ | |
| | **GP** | Forward | 111 | TGGGCTGAAAACTGCTACAATC | **45** |
| | | Reverse | | CTTTGTGCACATACCGGCAC | |
| | | Probe | | FAM-CTACCAGCA-ZEN-GCGCCAGACGG-IABkFQ | |
| | **VP30** | Forward | 109 | GCACCCAAGGACTCGCGCTT | **3** |
| | | Reverse | | TCGCCCAGTGTTCTGCCGTC | |
| | | Probe | | FAM-TCCAACGGC-ZEN-TGATGATTTCCAGCA-IABkFQ | |
| | **VP40** | Forward | 161 | ATCGAATCCACTCAGGCCAAT | **46** |
| | | Reverse | | CGACACCTAGAGGAAGCCAAA | |
| | | Probe | | FAM-AATGTCATA-ZEN-TCGGGCCCCAAAGTGC-IABkFQ | |
| Human | **GAPDH** | Forward | 143 | ACATCGCTCAGACACCATG | |
| | | Reverse | | TGTAGTTGAGGTCAATGAAGGG | |
| | | Probe | | Cy5-AAGGTCGGAGTCAACGGATTTGGTC-IAbRQSp | |

**Supplementary Table 2. Primers and probe sets used for detecting EBOV RNA genome by RT-qPCR.**

Primers and probes were constructed for targeting mRNA of each gene based on the indicated references and purchased form IDT after synthesizing by them. Human GAPDH was used as a housekeeping gene. Both reporter dye and quencher were attached to 5' and 3' of probes, respectively. ZEN quencher was attached in the middle od probes as described. ZEBOV = Zaire Ebola virus.
